# New classification of intrinsic disorder in the Human proteome

**DOI:** 10.1101/446351

**Authors:** Antonio Deiana, Sergio Forcelloni, Alessandro Porrello, Andrea Giansanti

## Abstract

We propose a new, sequence-only, classification of intrinsically disordered human proteins which is based on two parameters: dr, the percentage of disordered residues, and Ld, the length of the longest disordered segment in the sequence. Depending on dr and Ld, we distinguish five variants: i) *ordered proteins* (ORDs); ii) *not disordered proteins* (NDPsj; (iii) *proteins with intrinsically disordered regions* (PDRs); iv) *intrinsically disordered proteins* (IDPs) and v) *proteins with fragmented* disorder (FRAGs). PDRs have been considered in the general category of intrinsically disordered proteins for a long time. We show that PDRs are closer to globular, ordered proteins (ORDs and NDPs) than to disordered ones (IDPs), both in amino acid composition and functionally. Moreover, NDPs and PDRs are uniformly spread over several functional protein classes, whereas IDPs are concentrated only on two, namely *nucleic acid binding proteins* and *transcription factors*, which are just a subset of the functions that are commonly associated with protein intrinsic disorder. As a conclusion, PDRs and IDPs should be considered, in future classifications, as distinct variants of disordered proteins, with different physical-chemical properties and functional spectra.

## INTRODUCTION

In the last two decades, evidence and the concept of natively unfolded proteins, which are structurally disordered but functional, emerged as a prominent and influential theme in protein science^1-3^. The idea of an intrinsically disordered (natively unfolded, structurally heterogeneous) protein is that of a polypeptide which samples broad distributions of conformational parameters (e.g., gyration radii^4,5^). The opposite view is that of a globular (ordered) protein, such as a solid-like polymer in which diffusive motions are quite limited and which is, thus, prone to be folded into a three-dimensional (3D) structure, where each atom has a well-defined average equilibrium position. Many have used to call this new trend in protein science as the shift of a paradigm^6^. Despite the absence of a well-defined three-dimensional conformation, intrinsically disordered proteins are involved in important cellular functions, such as regulation, signaling^7-12^, alternative splicing^13-15^, as well as in the development of many diseases^16-18^, including cancer^19^, cardio-vascular^20^, and neurodegenerative diseases^21^. Many tumor suppressors have at least a large unstructured region, perhaps the best-known examples being p53 and BRCA1. Xie et al. in a series of three papers have described the biological functions of proteins with long disordered segments^22-24^. Furthermore, natively unfolded proteins are often the hubs of protein-protein interaction networks^25^.

Nowadays, after almost two decades of thorough experimental work (based on NMR and many other techniques: circular dichroism, fluorescence spectroscopy, vibrational CD spectroscopy, Raman spectroscopy, SAXS, to name just a few^26,27^) and an exponential growth of computational studies^17^, the field of intrinsically disordered proteins is mature both in pure and applied protein science. Nevertheless, while a remarkable set of consistent observations has emerged^28^, the mechanisms and functional spectra of natively unfolded proteins are still to be clearly established.

It is then of fundamental importance to have a reliable and fine-tuned classification of protein intrinsic disorder.

The focus of this work is on the distinction between two variants of proteins containing at least a long disordered segment (PDRs and IDPs), which are structurally and functionally distinct. The distinction between fully unstructured proteins (IDPs) and proteins which contain long disordered, unstructured regions (PDRs) is not new^2,29,30^; we make this distinction fine-tuned and quantitative, taking the human proteome as a case of study. Following the current view, an intrinsically disordered protein (IDP) is a protein that lacks a three-dimensional structure in a long portion of its polypeptide chain. IDPs do not interact with substrates through a lock-and-key mechanism but can interact with many substrates with high specificity and low-affinity^6,31^. The existence of mostly folded proteins but with intrinsically disordered regions (IDR) has been extensively discussed in the literature^30,32,33^. These latter proteins do keep a tertiary structure and therefore it is possible that they still interact with substrates through a lock-and-key mechanism. The presence of unstructured loops or domains (e.g. linear motifs, and Molecular Recognition Features - MoRFs) can be important for these proteins to target low affinity substrates, enlarging the overall repertoire of protein interactions^11^.

Many studies about classification and functional roles of disordered proteins are based on either the length of disordered regions in the sequence or the fraction of residues that is shown or predicted to be disordered^11^. It has been recently observed that proteins with at least one disordered segment longer than 30 amino acids have a higher turn-over with respect to other proteins in two cases: i) the segment is at least 30 amino acids long and it is located in the N or C terminus of the protein sequence; or ii) the segment is at least 40 amino acids long^11^. Moreover, about 33% of proteins from Eukaryotes have a long disordered segment (>30 amino acids)^30,34^; 44% of human proteins contain long disordered segments with more than 30 amino acids and 24% of them have at least 30% of residues predicted to be disordered^11^. Consequently, many studies in the field have identified IDPs as proteins with long disordered segments, i.e., segments of at least 30 consecutive disordered amino acids^35-40^. However, the presence of just one long disordered segment does not imply that a protein is intrinsically disordered, i.e., that it lacks a stable overall 3D conformation. Our distinction is based precisely on this point: the difference between IDPs and PDRs should depend on the relative proportion of the unstructured regions with respect to the rest of protein. It has indeed been shown that also proteins with a percentage dr of disordered residues larger than 30% have a higher turn-over in the cell^41^. Therefore, both length of disordered segments in a sequence and overall percentage of disorder are important parameters to distinguish variants of intrinsic disorder. This observation suggests the criterion that we propose in this study: a protein is intrinsically disordered (IDP) if it has at least one disordered segment of length Ld > 30 and more than 30% of its amino acids predicted as disordered. This criterion is indeed not completely new^15,26,42^. By contrast, we label here as PDRs proteins with at least one disordered segment of length Ld > 30, but with a percentage of disorder dr lower than 30%. We find here, in according to previous estimates^15^, that more than 50% of the human proteome contains long disordered regions (26% PDRs and 36% IDPs). Moreover, we found that PDRs have an amino acid composition which is more similar to those of ordered (ORDs) or not disordered proteins (NDPs) than to those of intrinsically disordered ones (IDPs), that have a high percentage of disorder promoting amino acids. Moreover, PDRs result also as more similar to ORDs and NDPs than to IDPs, on the basis of ontology and functional spectra. We concluded that PDRs and IDPs are distinct variants of disordered proteins and should be considered separately. This finding is clearly noteworthy, because separating PDRs from disordered proteins has the potential to improve the classification of the human proteome and clarify the role of disordered proteins in the cellular processes as well as in the emergence of diseases.

## EXPERIMENTAL SECTION

### Dataset of protein sequences

We downloaded the human proteome from the UniProt-SwissProt^43^ database (http://www.uniprot.org/uniprot), release of September 2017. We selected human proteins by searching for reviewed proteins belonging to the organism Homo Sapiens (Organism ID: 9606, Proteome ID: UP000005640). Out of 20177 protein sequences, 60 were discarded because contained atypical amino acid symbols (B, U, Z, X), thus generating a list with 20117 protein sequences.

### Disorder prediction

Each residue in the sequences of human proteins has been classified either as ordered or disordered using MobiDB^44^ (http://mobidb.bio.unipd.it). We have used MobiDB because it is a consensus database that combines experimental and curated data (expecially from X-ray crystallography, NMR and Cryo-EM), indirect sources of information on the disordered state of residues and disorder predictions for all UniProt entries using these tools: IUPred-short, IUPred-long, GlobPlot, DisEMBL-465, DisEMBL-HL, Espritz-DisProt, Espritz-NMR & Espritz-X-ray. The use of these predictors enables MobiDB to provide disorder annotations for every protein, even when no curated or indirect data is available.

### Classification of disordered proteins in the human proteome

To partition the human proteome into variants of disorder, we used two parameters: dr, the percentage of disordered residues in the whole sequence, and Ld, the length of the longest disordered domain in the sequence. We distinguish five variants of protein sequences, namely:

i. *Ordered proteins* (ORDs), that do not have disordered segments longer than 30 residues (Ld < 30) nor more than 10% of disordered residues (dr < 10%);
ii. *not disordered proteins* (NDPs), that do not have disordered segments longer than 30 residues (Ld < 30), with more than 10% but less than 30% of disordered residues (10% ≤ dr < 30%);
iii. *proteins with intrinsically disordered regions* (PDRs), that have at least one disordered domain longer than 30 residues (Ld ≥ 30) and are disordered in less than 30% of their residues (dr < 30%); iv) proteins that are *intrinsically disordered* (IDPs), that have at least one disordered segment longer than 30 residues (Ld ≥ 30) and that are disordered in more than 30% of their residues (dr ≥ 30%);
iv. *proteins with fragmented disorder* (FRAGs), that do not have a disordered fragment longer than 30 residues (Ld < 30) and that, nevertheless, have at least 30% of their residues predicted as disordered (dr ≥ 30%).

Following this distinction, ORDs and NDPs are, clearly, proteins with a limited number of disordered residues and absence of disordered domains (Ld < 30). We considered ORDs in order to represent a variant of completely ordered proteins with dr < 10%. PDRs, unlike NDPs, are proteins characterized by the presence of disordered domains (Ld ≥ 30) in structures that may well be globally folded. IDPs intended to be the true intrinsically disordered proteins, which not only embed disordered domains, but are also disordered in a relevant percentage of their residues. FRAGs, are a small number of proteins characterized by highly distributed disordered residues along the entire sequence.

### Euclidean distance between amino acid compositions

To represent differences in amino acid compositions 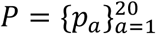 and 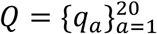, we used the Euclidean distance (E(P,Q)) which is formally defined as

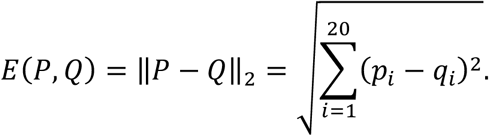

The Euclidean distance, at variance with other separations measures (e.g. Kullback-Leibler divergence), satisfies the mathematical properties of a distance function, i.e. non-negativity (being null between *P* and *Q* if and only if *P* = *Q*), symmetry, and triangle inequality and originates symmetric distance matrices.

The statistical significance of the observed distances *d_α,β_* between the amino acidic compositions of variant 1 and 2 in the distance matrix in Table 2 has been evaluated using a simple randomization test in which the *n_α_* sequences from the variant 1 and the *nβ* sequences from variant 2 are appended, renumbered from 1 to *n_α_ + n_β_*, and repeatedly shuffled to produce, at each iteration *i* two sets of *n_α_* and *n_β_* sequences whose amino acidic compositions are evaluated as *P_i_* and *Q_i_*. The null hypothesis to be tested is that amino acid compositions (*P* and *Q*) of the pair variants came from the same distribution and the p-value was estimated as

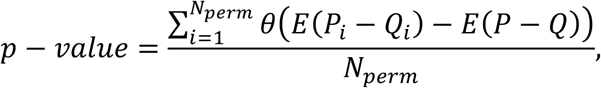

where *θ* is the Heaviside step function, and *N_perm_* is the number of shuffling iterations *(N_perm_ =* 10000). This p-value has the straightforward meaning of the probability of getting, from random sampling two given variants, a compositional distance that is bigger than the observed one^45^.

### Functional screening and ontologies

The four variants of proteins (ORDs, NDPs, PDRs, IDPs and FRAGs) were related to several protein classes and ontologies, following the annotations of the Gene Ontology Consortium^46^, according to their molecular functions, biological processes and cellular components. We used the PANTHER (Protein Annotation Through Evolutionary Relationship)^47-49^ classification system (http://www.pantherdb.org). To evaluate the functional profiles of each variant we estimated the conditional probability *P* (*protein class* | *variant*), by computing the number of proteins of that variant in each PANTHER class divided by the total number of proteins in the specified variant. To evaluate how a PANTHER protein class is enriched in a variant of disorder we estimated the conditional probability *P* (*variant* | *protein class*), by computing the number of proteins in the class which belong to each one of the protein variants: ORDs, NDPs, PDRs, IDPs and FRAGs; then, these numbers were divided by the total number of proteins belonging to the specific class.

### Binomial statistical test of enrichment in variants of disorder

To assess the enrichment of a PANTHER protein class in a specific variant (or group of variants) again another a statistical test based on the binomial distribution was used^50^. Suppose that the proteins belonging to a class are separated into two groups: A and B. The occurrences of proteins of type A (e.g. not disordered) and the occurrences of proteins of type B (e.g. disordered), divided by the total number of proteins in the class are estimates of the probabilities of type A or B to belong to that class. In the test, the null hypothesis (H_0_) is that neither of the two groups outnumbers the other; in other words, proteins in the two groups are sampled from the same general population, and thus the probability of observing a protein in one of the two groups is the same as in the other group. Through the binomial test the observed difference is tested and, if the evaluated p-value is less than the threshold of 0.0001, then the null hypothesis is rejected and the observed difference is considered statistically significant.

### Analysis of interaction interfaces in variants of disorder

Considering the different variants of disorder (ORDs, NDPs, PDRs, IDPs, and FRAGs) as a case of study, we studied the properties of interaction interfaces. The interfacial regions are selected from Interactome3D^51^ (https://interactome3d.irbbarcelona.org), which is a web service for the structural annotation of protein-protein interaction networks. For each protein, we calculated the fraction of complexed residues which are predicted disordered by MobiDB, with respect to the total number of complexed residues:

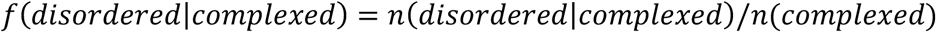

The total length of the complexed regions was used as normalization factor, in order to avoid bias due to the percentage of disordered residues that characterize the variants of disorder.

## RESULTS AND DISCUSSION

### Variants of disorder in the human proteome

As specified in the Materials and Methods, 20117 sequences of human proteins were extracted from UniProt/SwissProt and selected from the MobiDB database. Only 15751 of them (79%) were functionally annotated by the PANTHER classification system. These sequences were partitioned into five variants. In Table 1 we show the coverage of the variants of disorder in the human proteome. Interestingly, if one considers PDRs as disordered proteins, together with IDPs and FRAGs, then the percentage of estimated disordered in the human proteome is about 55%, consistently with previous global estimates^15^. Conversely, if one considers PDRs as a variant of ordered proteins and puts them together with ordered (ORDs) and not disordered proteins (NDPs), then the amount of disorder in the human proteome is reduced to about 28%.

**Table 1:** Number of ORD, NDP, PDR, IDP, and FRAG proteins in the human proteome.

| Variant | MobiDB | % | PANTHER | % |
| --- | --- | --- | --- | --- |
| ORD | 6031 | 30.1 | 5062 | 32.1 |
| NDP | 2975 | 14.8 | 2161 | 13.7 |
| PDR | 4558 | 22.8 | 4186 | 26.6 |
| IDP | 6051 | 30.2 | 4203 | 26.7 |
| FRAG | 415 | 2.1 | 139 | 0.9 |
| Totals | 20030 | 100 | 15751 | 100 |

Number of human proteins belonging to the different variants and corresponding coverage in the MobiDB and PANTHER databases. Proteins with more than 30% of predicted disorder (PDRs, IDPs, and FRAGs) constitute 55% of the human proteome. Not disordered proteins (ORDs and NDPs) make the remaining 45%. Note that if PDRs are considered ordered then the estimated fraction of disordered proteins in the human proteome drops to 28%.

Therefore, the classification of PDRs either as ordered or disordered is *quite relevant to estimate the overall percentage of disordered proteins in the human proteome.* FRAGs, characterized by a sparse distribution of disorder, are represented by a few cases worth being investigated, in future studies, both as a category and as single cases; most of these proteins are rich in cysteine residues which stabilize their structure through the formation of disulfide bridges.

### Variants of disorder: amino acid compositions

To summarize the differences in amino acid compositions of the protein variants, we calculated the Euclidean distance between each pair of distributions and verified the significance of these distances performing a randomization test with 10000 iterations (see Materials and Methods). Analyzing the values of distances collected in Table 2, PDRs are less distant to ORDs (0.025) and NDPs (0.013) than to IDPs (0.049); in other words, the distance between PDRs and IDPs is significantly about four times larger than the distance between PDRs and NDPs.

**Table 2.** Euclidean distances between amino acid compositions of the variants.

|  | ORDs | NDPs | PDRs | IDPs | FRAGs |
| --- | --- | --- | --- | --- | --- |
| ORDs | 0,0 | 0,025 | 0,025 | 0,071 | 0,054 |
| NDPs | - | 0,0 | 0,013 | 0,051 | 0,032 |
| PDRs | - | - | 0,0 | 0,049 | 0,035 |
| IDPs | - | - | - | 0,0 | 0,030 |
| FRAGs | - | - | - | - | 0,0 |

Each entry of this symmetric matrix represents the Euclidean distance between the amino acid distributions of the corresponding variants reported in the first raw and in the first column. From the point of view of the amino acid compositions, PDRs are more similar to ORDs and NDPs than to IDPs and FRAGs. Each distance is significant with respect to the random case (p-value<0.0001).

These first observations show that PDRs and IDPs are characterized by different amino acid repertoires that should be reflected in their different chemical-physical properties.

### Functional spectra of ORD, NDPs, PDRs, IDPs, and FRAGs amongst PANTHER protein classes

For each variant of disorder, a unitary vector whose components are the conditional probabilities *P*(*protem-class*|*variant*) has been evaluated over the 26 protein classes of the PANTHER classification system. These conditional probabilities measure the propensity of each variant of disorder to fall into the different PANTHER protein classes and synthetically represent the functional spectrum of each variant (Fig. 1).

**Figure 1.**
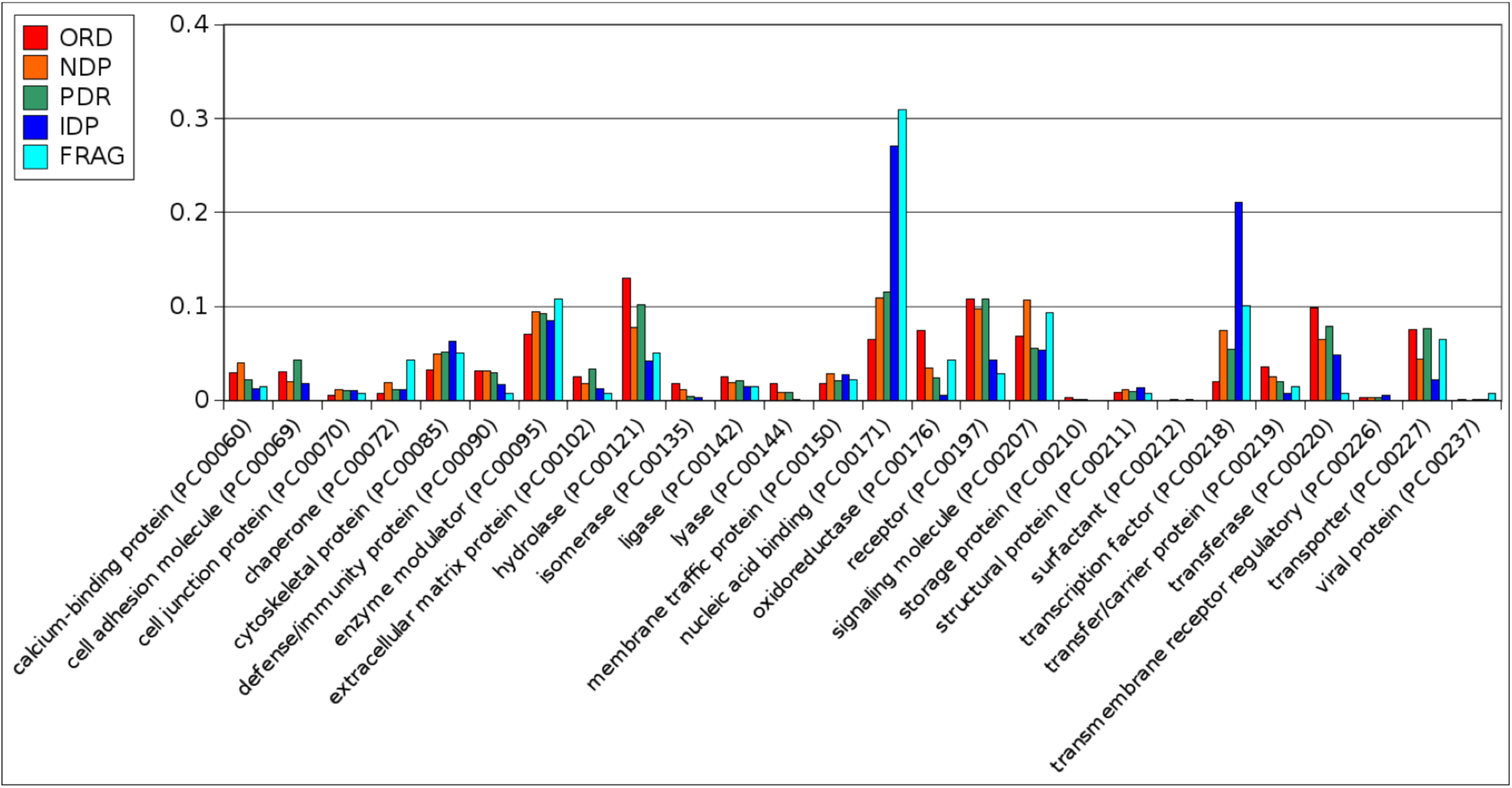
Functional spectra of variants of disorder in the human proteome. Each colored histogram represents the functional spectrum of a variant of disorder, as a normalized vector of frequencies that estimate the conditional probabilities *P*(*protein-class*|*variant*). Each component of these vectors estimates the relative probability of a variant to belong to different protein classes.

A quantitative measure of the functional similarities between variants is given by the matrix encoding Euclidean distances between functional spectra (Table 3).

**Table 3.** Functional distance matrix between variants of disorder in the human proteome.

|  | ORDs | NDPs | PDRs | IDPs | FRAGs |
| --- | --- | --- | --- | --- | --- |
| ORDs | 0.0 | 0.12 | 0.10 | 0.32 | 0.31 |
| NDPs | - | 0.0 | 0.08 | 0.23 | 0.23 |
| PDRs | - | - | 0.0 | 0.25 | 0.24 |
| IDPs | - | - | - | 0.0 | 0.15 |
| FRAGs | - | - | - | - | 0.0 |

The entries of this symmetric matrix were computed as the modulus of the difference between the functional spectra *P*(*protein class*|*variant*) associated with each variant of disorder (i.e. Euclidean distances). Refer to Fig. 1.

The relevant observation is that PDRs are less distant, functionally, from ORDs (0.10) and NDPs (0.12) than from IDPs (0.25) and FRAGs (0.24). In other words, the functional spectrum of PDRs is more overlapped with those of ORDs and NDPs than with those of IDPs and FRAGs. This observation clearly supports separating PDRs from IDPs, at variance with the usual classification of PDRs as IDPs.

### Enrichment of PANTHER protein-classes in variants of disorder

To complete the information given in the previous section, it is interesting to evaluate how each PANTHER protein-class is enriched in one or more of the five variants. We thus computed, among the proteins that belong to a specific class, the fraction of each variant of disorder. This is a way of estimating *P*(*variant*|*protein-class*) (Fig. 2). Visually, several classes seem to be dominated by a variant which is undoubtedly more frequent than the others (e.g. class PC00135 is enriched in ORDs, whereas class PC00211 in IDPs). In other classes, the situation is less clear (e.g. in class PC00095).

**Figure 2.**
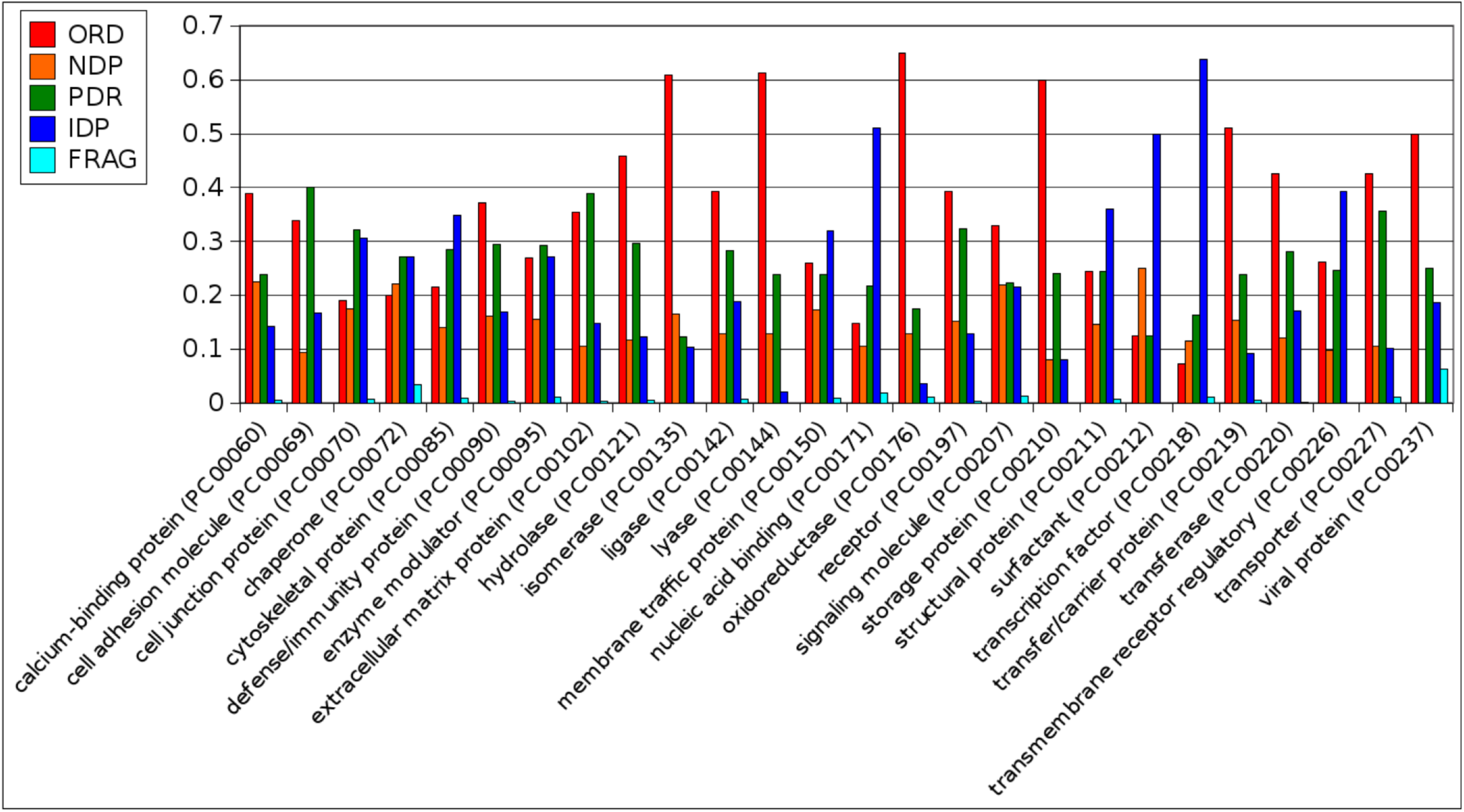
Relative occurrence of ORDs, NDPs, PDRs, IDPs and FRAGs proteins in PANTHER protein classes. These histograms, complement those in Fig. 1 and represent the conditional probabilities *P*(*variant*|*protein class*); the proteins in each one of the 26 PANTHER protein-classes are partitioned into the five variants of disorder; the normalization is over each protein class. In these histograms, FRAGs are associated with the shortest column, due to their limited number (Table 1).

To be more precise in deciding about enrichments we used a binomial test for assessing the difference between the frequency of the variant of disorder in the PANTHER protein-class and the frequency of the same variant in the whole human proteome^50^. Because, in each case, we performed several statistical tests, the p-values have been corrected using the method of Bonferroni.

In Table 4, we have collected the PANTHER protein classes that are enriched in a particular variant of disorder in a statistically significant way (p-value<0.0001).

**Table 4.** PANTHER protein classes that are enriched in variants of disorder.

| ORDs | NDPs | PDRs | IDPs |
| --- | --- | --- | --- |
| *Calcium binding proteins* (PC00060)<br>*Hydrolases* (PC00121)<br>*Isomerases* (PC00135)<br>*Ligase* (PC00142)<br>*Lyases* (PC00144)<br>*Oxidoreductases* (PC00176)<br>*Receptor* (PC00197)<br>*Transfer/carrier proteins* (PC00219)<br>*Transferases* (PC00220)<br>*Transporter proteins* (PC00227) | *Calcium binding proteins* (PC00060)<br>*Signaling molecule* (PC00207) | *Cell adhesion molecules* (PC00069)<br>*Cytoskeletal proteins* (PC00085)<br>*Enzyme modulator* (PC00095)<br>*Extracellular matrix proteins* (PC00102)<br>*Hydrolases* (PC00121)<br>*Receptor* (PC00197)<br>*Transferases* (PC00220)<br>*Transporter proteins* (PC00227) | *Nucleic acid binding* (PC00171)<br>*Transcription factors* (PC00218) |

This table collects PANTHER protein classes which are enriched in a statistically significant way (p-value<0.0001) in one of the four variants: ORDs, NDPs, PDRs, and IDPs. FRAGs are not shown because were deemed not statistically relevant, due to their limited number (Table 1).

Interestingly, IDPs clearly dominate among *nucleic acid binding* (PC00171) and *transcription factors* (PC00218). ORDs are over-represented in *calcium binding proteins* (PC00060), *transporter proteins* (PC00090), *transfer/carrier proteins* (PC00219), *receptor* (PC00197) and, as expected, among enzymes: *hydrolases* (PC00121), *isomerases* (PC00135), *ligase* (PC00142), *lyases* (PC00144), *oxidoreductases* (PC00176), and *transferases* (PC00220). We note that some protein classes dominated by PDRs refer to structural proteins such as *extracellular structural proteins* (PC00211), *cell adhesion molecules* (PC00069), and *cytoskeletal proteins* (PC0085), that, interestingly, are usually rigidly structured proteins. On the other hand, looking at Table 5, we clearly observe a strong depletion of the IDP variant in many PANTHER protein classes, thus decentralizing and specializing the role of mostly unstructured proteins in the cell.

**Table 5.**
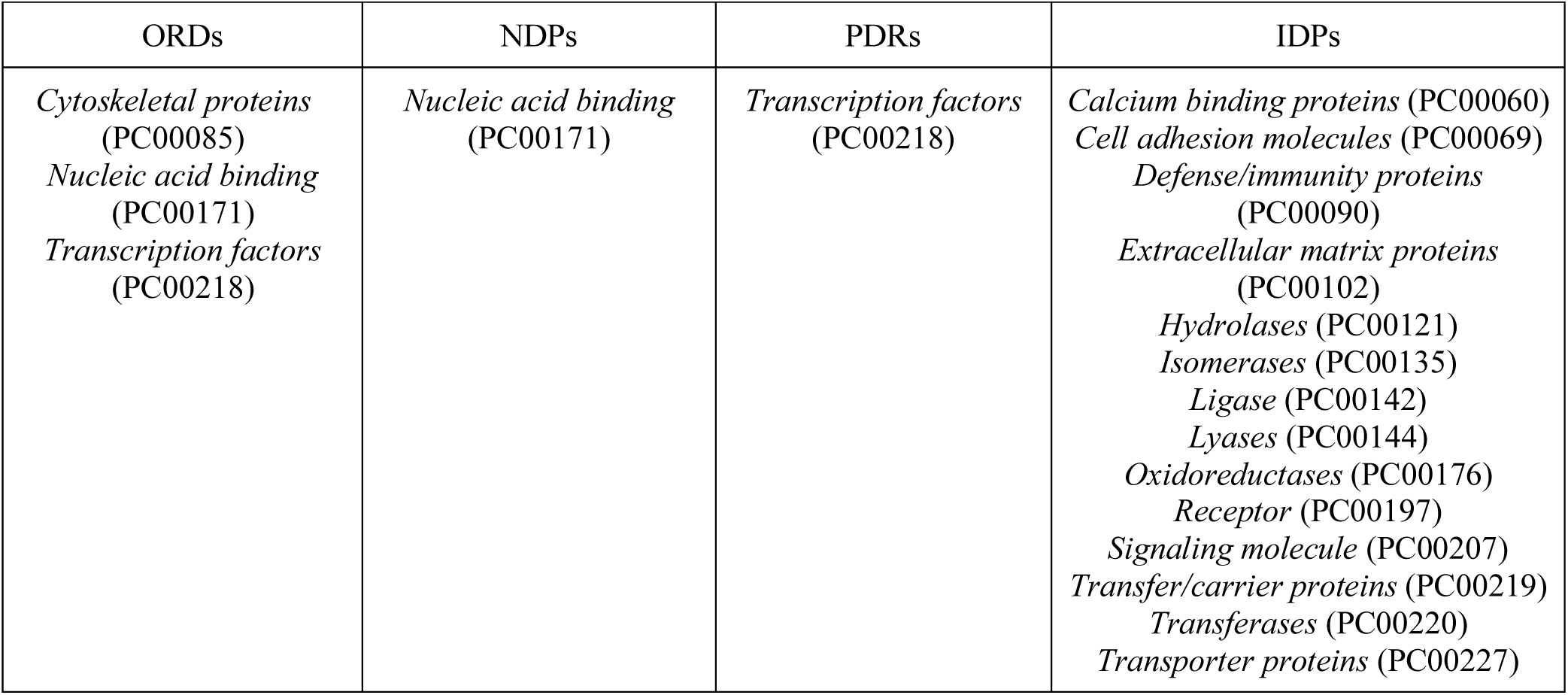
PANTHER protein classes that are depleted in variants of disorder.

This table collects PANTHER protein classes which are depleted in a statistically significant way (p-value<0.0001) in one of the four variants: ORDs, NDPs, PDRs, and IDPs. FRAGs are not shown because were deemed not statistically relevant, due to their limited number (Table 1).

The main purpose of this work is to establish whether PDRs should be considered disordered or not. To shed some light on this subject, we performed tests of enrichment in two cases. In the first, we consider PDRs as ordered, and put them in the same group as ORDs and NDPs, as suggested by the observation that their functional spectra are closer; in the second, we put PDRs in the same group of IDPs and FRAGs, as also done by other authors^35-40^. The statistical method used to assess the enrichment of PANTHER classes in proteins of the disordered group over the other is the binomial test (see Materials and Methods); in this case, a Bonferroni correction is not required because the comparison is made between two groups of proteins (ordered and disordered). It is clear from Table 6 that considering PDRs as ordered or disordered leads to a profound difference in the role of disorder within cellular processes. In particular, if PDRs are considered disordered, as well as IDPs and FRAGs, we obtain results in agreement with those previously reported in the literature^7-15,22-24^. Conversely, if PDRs are seen as a specific variant of ordered proteins, the landscape about protein disorder changes dramatically. Indeed, in the latter case, the PANTHER protein classes enriched in disordered proteins are only *nucleic acid binding* (PC00171), and *transcription factors* (PC00218).

**Table 6.** Different features associated with disordered proteins considering the PDRs as ordered or disordered.

| **1<sup>st</sup> Test: PDRs as ordered** | **(ORDs+NDs+PDRs) vs (IDPs+FRAGs)** |
| --- | --- |
| PANTHER class | p-value |
| *Nucleic acid binding* (PC00171) | $1.4 \times 10^{-8}$ |
| *Transcription factors* (PC00218) | $< 10^{-16}$ |
| **2<sup>nd</sup> Test: PDRs as disordered** | **(ORDs+NDs) vs (PDRs+IDPs+FRAGs)** |
| *Cell adhesion molecules* (PC00069) | $9 \times 10^{-8}$ |
| *Cell junction proteins* (PC00070) | $< 10^{-16}$ |
| *Chaperones* (PC00072) | $3 \times 10^{-5}$ |
| *Cytoskeletal proteins* (PC00085) | $< 10^{-16}$ |
| *Enzyme modulator* (PC00095) | $< 10^{-16}$ |
| *Membrane traffic proteins* (PC00150) | $3 \times 10^{-7}$ |
| *Nucleic acid binding* (PC00171) | $< 10^{-16}$ |
| *Structural proteins* (PC00211) | $2 \times 10^{-8}$ |
| *Transcription factors* (PC00218) | $< 10^{-16}$ |
| *Transmembrane receptor regulatory* (PC00226) | $1.2 \times 10^{-5}$ |

PANTHER protein classes that are enriched in disordered variants in two statistical tests: in the first one, PDRs are considered as a variant of ordered proteins; in the second, they are considered as disordered, together with IDPs and FRAGs. Note the remarkable change in the functional spectrum associated to disorder induced by the shift in the classification of PDRs.

### Fraction of disordered residues in interaction interfaces

To further investigate the differences between variants of disorder, we considered how the different variants of disorder are present in the molecular interfaces associated with protein-protein interaction. In particular, we have estimated the fraction of disordered residues that, in each variant, belong to interaction interfaces as structurally annotated in Interactome3D. The number of proteins in each variant for which we have informations on the complexes in Interactome3D are shown in Table 7. The average coverage is 29% and *no variant of proteins appears to be particularly over-represented in Interactome3D.*

**Table 7.** Coverage of each variant of disorder in Interactome3D.

| Variants of disorder | Number of complexed proteins in Interactome3D | Number of proteins in the variants (Table 1) | % complexed proteins in each variants |
| --- | --- | --- | --- |
| ORDs | 1698 | 6031 | 28 |
| NDs | 948 | 2975 | 32 |
| PDRs | 1495 | 4558 | 33 |
| IDPs | 1890 | 6051 | 30 |
| FRAGs | 90 | 415 | 22 |

The number (second column) and percentage (fourth column) of complexed proteins annotated in Interatome3D for each variant of disorder are shown. The coverage in Interactome3D is quite uniform among the five variants.

For each protein annotated in Interactome3D, we counted the number of residues used in the interaction with other partners (n(complexed)), and the number of residues in complex which are predicted to be disordered (n(disordered | complexed)) (Fig. 3).

**Figure 3.**
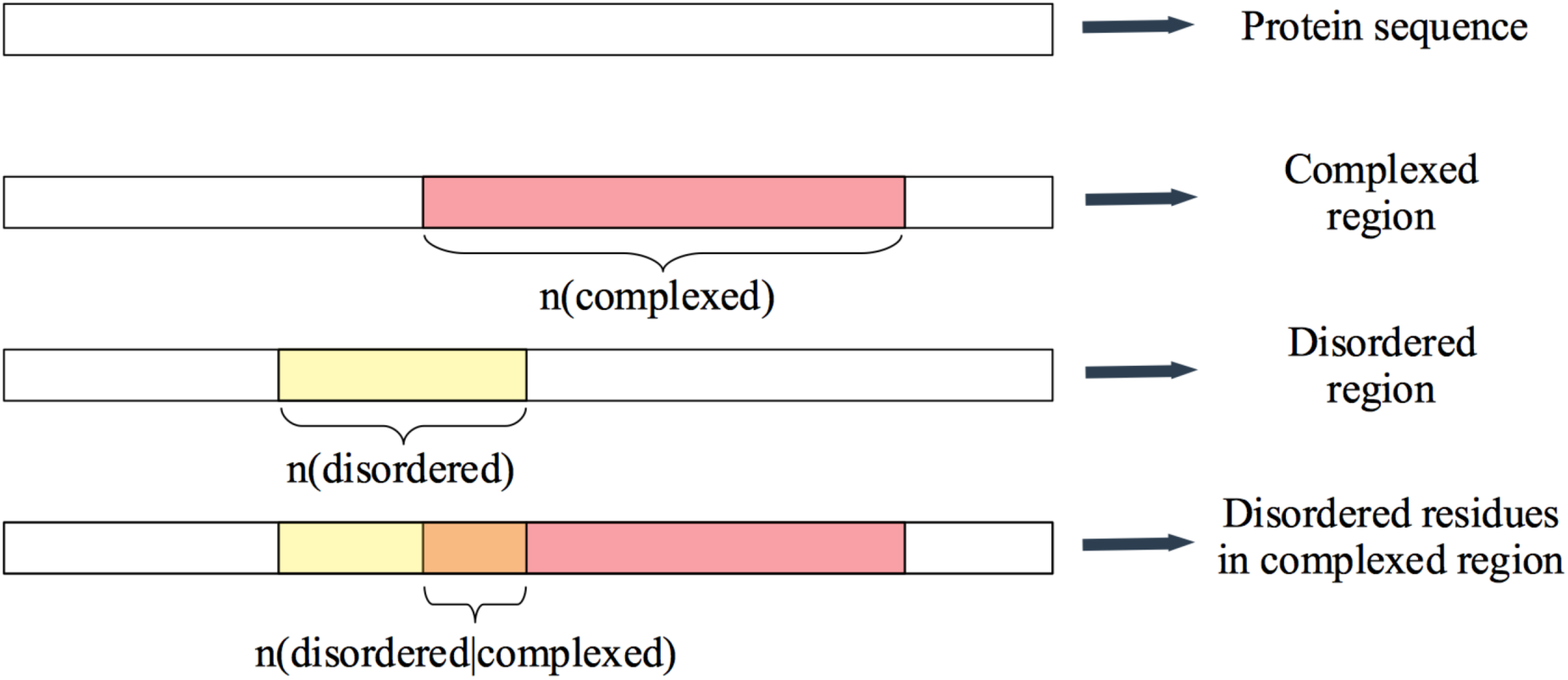
Example of a complexed and disordered region in a protein sequence. A schematic representation of the quantities n(complexed), n(disordered), n(disordered | complexed) is shown. In pink, we display the complexed region and, in yellow, the predicted disordered segment in the sequence; their intersection (in orange) represents the fraction of disordered amino acids in the interaction interface.

Specifically, for each protein covered in Interactome3D, we evaluated the fraction f(disordered | complexed) defined as the number of complexed residues predicted to be disordered with respect to the total length of the interaction interface (Materials and Methods). In Fig. 4 we report the mean value of f(disordered | complexed) for each variant of disorder.

**Figure 4.**
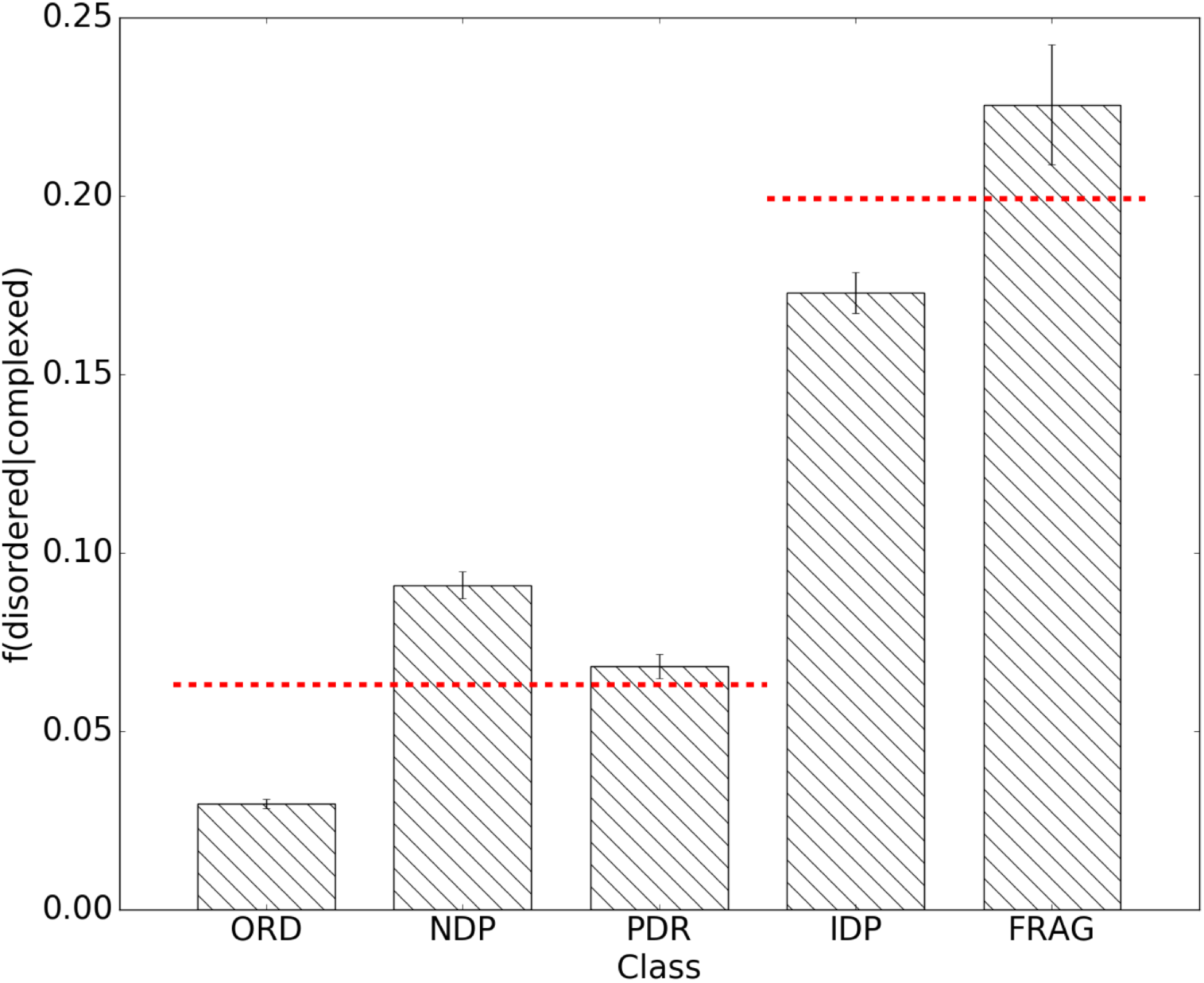
Different usage of disordered residues in interaction interfaces. For each variant, we show the mean values of f(disordered | complexed), the fraction of disordered residues engaged in interaction interface, as annotated in Interactome 3D. Error bars are standard deviations over each variant. Note that there are two separated mean levels (red broken lines).

Since these distributions are not normal, we used the Wilcoxon test with the Bonferroni correction to establish if the samples are pairwise statistically different. Indeed, we checked that f(disordered | complexed), the fraction of disordered residues engaged in interaction interfaces, has a different statistic in each variant (Fig. 4). Clearly f(disordered | complexed) is about 6% for the group {ORDs, NDPs, PDRs} and about 20% for the group made by IDPs and FRAGs. Also in this case IDPs are closer to ORDs and NDPs than to IDPs and FRAGs. PDRs and IDPs are then clearly characterized by a different propensity to recruit disordered residues in their interaction interfaces.

## CONCLUSIONS

In many papers in this field, a protein is considered disordered when it has a disordered region longer than 30 residues. We revisited this definition considering also the percentage of disordered residues with respect to the total length of the proteins. We define IDPs as proteins with Ld > 30 and dr > 30%. On the other hand, if a protein has a long disordered segment but less than 30% of disordered amino acids, we consider it a PDR. It is important to point out that PDRs and IDPs have both a long disordered segment; therefore, in a part of the current literature^35-40^, PDRs and IDPs have been considered a unique variant of intrinsic disorder. Clearly, if a protein has long disordered segments but, nevertheless, is mostly ordered in the rest of the sequence, it is quite hard to consider it a fully intrinsically disordered protein since, it has, in many cases, a well-defined three-dimensional structure. The focus of this work is on the distinction between PDRs and IDPs. IDPs have very long disordered segments that account for a large proportion of the total polypeptide chain, whereas PDRs are proteins with intrinsically disordered regions inserted as flexible “extensions” in structural context. Using a straightforward classification, based on the presence of long disordered domains (as often done) in conjunction with the evaluation of percentages of disordered residues, we found that 26% and 36% of human proteins are PDRs and IDPs, respectively (Table 1). Once one has separated PDRs as a category of proteins with localized disorder, the main question is the following: *are PDRs functionally and structurally closer to the ordered categories (ORDs and NDPs) than to disordered categories (IDPs and FRAGs)?* This study showed that PDRs are a variant of globular, structured, ordered proteins, as we shall further investigate with a survey of PDRs that are in the Protein Data Bank, revisiting previous works^52,53^. Interestingly, PDRs have an amino acid composition (Table 2) more similar to those of ORDs and NDPs than to that of intrinsically disordered proteins (IDPs), which are instead rich in disorder-promoting (hydrophilic and charged) amino acids and poor in order-promoting (hydrophobic, apolar, bulky) amino acids. This key observation points to the fact that, based on physical-chemical properties of their sequences, PDRs are more similar to folded than to unfolded proteins. Using the PANTHER database, we have also seen that PDRs have gene ontologies and functional profiles that are closer to those of non-disordered proteins than to those of IDPs (Table 3). As it is shown in Table 4, NDPs and PDRs are rather uniformly spread over different functional protein classes, whereas IDPs are particularly relevant for some specific functions, such as *nucleic acid binding* and *transcription factor.* Similarly, IDPs are under-represented in many PANTHER protein classes (Table 5) leading us to conjecture that unstructured proteins play a specialized role in the cell, at odds with what has been often stated until now.

Finally, PDRs and IDPs show a statistically significant difference in their tendency to interact with other proteins in complexes using different proportions of disordered residues at the interaction interfaces: PDRs tend to interact with other proteins by exploiting a smaller number of disordered residues (~6%) than intrinsically disordered variants IDPs and FRAGs (Fig. 4).

Taken together, our results suggest that the distinction between PDRs and IDPs is very important to correctly classify the human proteome. Therefore, these two classes should be regarded as two different and general protein variants, with both different physical-chemical properties and a different functional spectrum. This new classification could be very important in understanding functional aspects related to different types of proteins, the role of the intrinsic disorder in the cell and in the emergence of diseases.

Further investigations are surely needed. In particular, it could be useful to improve this classification and the differentiation between PDRs and IDPs, studying their separations as a function of the threshold of the parameters (Ld and dr). Moreover, it would be interesting to investigate the roles of each variant in protein-protein interaction networks.

## ABBREVIATIONS

ORD: ordered protein

NDP: not disordered protein

PDR: protein with disordered regions

IDP: intrinsically disordered protein

FRAG: intrinsically disordered protein with fragmented disorder

## SUPPORTING INFORMATIONS

Amino acidic distributions of variants of disorder.

